# Arabidopsis actin-binding protein WLIM2A links PAMP-triggered immunity and cytoskeletal organization

**DOI:** 10.1101/2023.12.13.571563

**Authors:** Prabhu Manickam, Aala A Abulfaraj, Hanna M Alhoraibi, Alaguraj Veluchamy, Marilia Almeida-Trapp, Heribert Hirt, Naganand Rayapuram

## Abstract

MAPKs are a family of highly conserved serine/threonine protein kinases that link upstream receptors to their downstream targets which can be localized in the cytoplasm or the nucleus. Pathogens produce pathogen-associated molecular patterns (PAMPs) that trigger the activation of MAPK cascades in plants. Phosphoproteomic analysis of PAMP-induced *Arabidopsis* plants led to the identification of several putative MAPK targets, WLIM2A. Here, we investigated the role of WLIM2A in plant immunity via a reverse-genetics approach generating *wlim2a* knockout lines using CRISPR-Cas9, as well as complementation and phosphosite mutated *WLIM2A* expression lines in the *wlim2a* background. The *wlim2a* lines were compromised in their response to *Pst* DC3000 but showed enhanced resistance to fungal infection by *Botrytis cinereae*. Transcriptome analyses revealed that immune hormone signaling and biosynthesis genes of salicylic acid (SA), jasmonic acid (JA), and ethylene (ET) are differentially regulated in the *wlim2a* knockout lines. Pathogen assays with *Pst DC3000* showed altered stomatal phenotypes in *wlim2a* mutants. Importantly, *WLIM2A* phosphomutants had opposing stomatal behaviour and resistance phenotypes in response to *Pst* DC3000 infection. Overall, these data show that phosphorylation of WLIM2A by MAPKs regulates *Arabidopsis* stomatal immunity.

## Introduction

PAMP-triggered immunity (PTI) refers to plant defense mechanisms triggered by PAMPs (Jones and Dangl et al. 2006; Underwood et al. 2012). Microbes have also developed host-pathogen interaction effector proteins, which are secreted into the host plant via the type-III secretion system (T3SS). The PTI pathway is inhibited by various effectors, resulting in effector-triggered susceptibility (ETS) (Jones et al., 2016). Plants have developed R genes that recognize pathogen effectors and trigger defensive responses via effector triggered immunity (ETI) (Dang and Jones et al. 2001 & 2006).

Numerous studies in the last ten years have shed light on how mitogen activated protein kinases (MAPKs) activate PAMP-triggered immunity during plant-pathogen interactions. MAPKs are well-known for their participation in numerous signaling pathways. Given the complexities of MAPK signaling, only a subset of MAPK targets has been discovered so far. We recently carried out a comparative phosphoproteomics analysis in WT and *mapk (mpk3, mpk4, mpk6)* mutants was carried out utilizing flg22 as a microbe-associated molecular pattern (MAMP). The search yielded 152 differentially phosphorylated peptides, 70 of which were identified as potential MAPK targets (Rayapuram et al., 2018). Of the 70 targets, At2g39900 (WLIM2A) was selected for further study because its role in plant immunity is unknown.

WLIM2A belongs to the family of LIM proteins, which are defined by having one to five LIM domains that associate with other domains, including homeodomains, catalytic domains, cytoskeleton binding domains, and protein binding domains (SH3 or LD domains) (Kadrmas and Beckerle et al. 2004; **Fig. S1C & S1D**). The LIM proteins are a eukaryotic protein family that is well studied in animals, but not much is known about plant LIM family proteins. Recently, it was shown that plant LIM proteins could play multiple roles in plant development, metabolism, and defense (Li et al., 2014; Xu et al. 2015; Li et al., 2015; Srivatsa and Verma et al. 2017; **Fig. S1A & B**). Plant LIM proteins are actin-bundling proteins (ABPs) whose LIM domains interact directly with actin filaments (Papuga et al., 2010). *Nicotiana tabacum* WLIM1 and *Lily* L1LIM1 have been shown to directly bind to actin filaments and crosslink actin filaments into actin bundles (Thomas et al., 2006; Wang et al., 2008). L1LIM1 promoted filamentous actin bundle assembly and protected F-actin filaments against latrunculin B-mediated depolymerization. The appearance of asterisk-shaped F-actin aggregation and the hyper bundle has been associated with a defective targeting of endomembrane trafficking and thereby impaired pollen tube elongation. Furthermore, a pleiotropic morphology is observed in pollen tubes overexpressing *L1LIM1*, with retarded pollen development, swollen tip, and multiple tubes emerging out of single pollen grains. These results suggest that L1LIM1 plays an important role in endomembrane trafficking by contributing to actin bundle formation or elongation of pollen tubes (Wang et al., 2008).

Recently, *Arabidopsis* LIM proteins were identified as RNA binding proteins (RBPs) in an mRNA interactome study (Reichel et al., 2016). Several studies suggest that the role of RBPs in RNA transport along the cytoskeleton facilitates mRNA localization (Bullock et al., 2011; Gagnon and Mowry et al., 2013), which suggests that *Arabidopsis* LIMs might also be involved in mRNA transport along actin filaments. In addition, actin function was also studied in the nucleus where it is involved in mRNA processing, export, and localization (Hofmann et al., 2009; Medoni et al., 2012). *LIM4* has been shown to bind to actin filaments and bundle them in response to Ca^2+^. Growing evidence points to the importance of the actin cytoskeleton in plant immunity. Actin filament abundance increases exponentially within minutes after receptor activation in *Arabidopsis* epidermal cells (Henty-Ridilla et al., 2013). This phenomenon is assumed to represent a unique early feature of PTI responses. The targeted transport of defense chemicals to the infection site, organelle rearrangements, and ligand-induced endocytosis of receptors are all examples of defense responses that require actin remodeling. Furthermore, callose deposition, apoplastic ROS production, and transcriptional reprogramming of defense genes are all significantly reduced when the host actin cytoskeleton is disturbed. Thus, the actin cytoskeleton and associated cellular mechanisms assist the organization of intracellular and apoplastic defenses in host plants. When the actin cytoskeleton is disrupted, both pathogenic and nonpathogenic bacteria are more likely to affect plants. But so far little is understood about the precise functions of the actin cytoskeleton in host defense.

The actin array in guard cells undergoes dynamic remodeling, which is required for effective stomatal closure. The actin filaments in the closed stomata reorganize from a radial array to a randomly organized network, and then to a longitudinal alignment. Disrupting this reorganization via genetic or pharmacological methods all results in impaired stomatal closure. The dynamic behaviors of individual actin filaments between guard cells of stomata during the open and closed stages were compared by Li et al. (2019). It is still uncertain what the first dynamic actin filament events are that may have caused the change to the actin structure connected to stomatal closure. Most research on actin organization in guard cells so far has been done under non-biotic stress conditions. Shimono et al. (2016) investigated the modifications to actin architecture that take place during the pathogen- and MAMP-induced stomatal closure. Their findings imply that guard cell actin array patterns during immunity are comparable to those during diurnal cycling. Further research is needed to understand the details of actin dynamics during stomatal defense and the underlying molecular pathways.

In this study, we report that WLIM2A interacts and is phosphorylated by three immune MAPKs (MPK3, MPK4, and MPK6). We demonstrate that *Arabidopsis wlim2a-1* and *wlim2a-2* knockout mutant lines have a reduced basal immune function and are more susceptible to the bacterial pathogen *P. syringae pv. tomato* DC3000 *(Pst)*. Transcriptome data analysis showed that *WLIM2A* regulates the transcription of defense-related genes and hormones. Furthermore, using pathogen assays, we clearly demonstrate an important regulatory role of WLIM2A phosphorylation during plant immunity. In response to *Pst* infection, we observed that in contrast to WT and *wlim2a-2 phosphomimicking mutant* plants, stomata in *wlim2a and wlim2a-1* phosphodead mutants are compromised in stomatal closure during pathogen attack, indicating that MAPK-directed WLIM2A phosphorylation plays a role in stomatal immunity.

## Materials and Methods

### CRISPR Cas9 mutant generation

Twenty nucleotide-long gRNAs specific to the target gene WLIM2A were created using the CRISPR-P 1.0 method in order to produce a *wlim2a* knockout mutant *Arabidopsis* line (Wang ZP et al., 2015). For each gene, we selected two gRNA targets with the lowest off-target scores. As previously described (Wang ZP et al., 2015), the gRNA cassettes were cloned into a mcherry pHSE401 vector (**Table. 1;** Xing et al., 2014) containing an egg cell-specific promoter, as described previously (Wang ZP et al., 2015). These plants were then transformed into *A. thaliana* Col-0 plants using the agro-floral dip technique (**Fig. S2**).

**Table 1.**
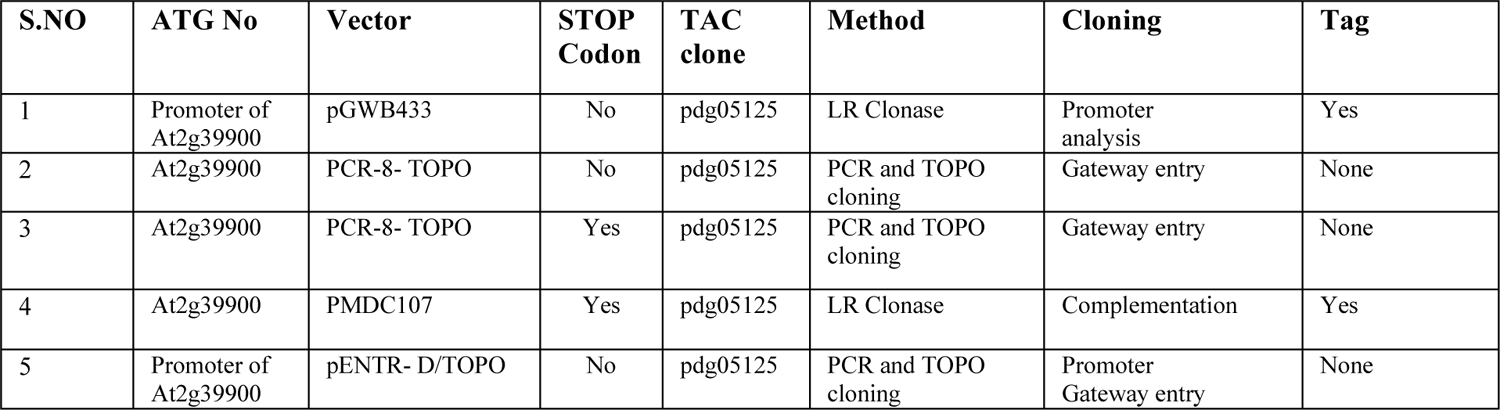
List of Constructs and Vectors.

### Quantitative Real-Time PCR

For MAMP induced defense gene expression, 14-day-old *Arabidopsis* WT and mutant seedlings grown on ½ MS agar were transferred onto liquid ½ MS and cultured overnight before flg22 treatment. Flagellin 22 (flg22) peptide was dissolved in double-distilled H_2_O to obtain a stock solution of 1 mM, then further diluted to inoculate at a concentration of 1 µM. Seedlings were treated with either 1 µM flg22 or water (mock) for 1 hour. WT and mutant seedlings samples (100 mg) were homogenized in liquid nitrogen, total RNA was extracted using NucleoSpin Plant RNA kits, and cDNA was reverse transcribed from 1 µg of total RNA using SuperScript III First-Strand Synthesis SupermixKit (Invitrogen), and the transcript levels of genes were assessed by qRT-PCR using the PTI marker gene primers (**Table. 2**). Quantitative RT-PCR was performed using SsoAdvanced Universal SYBR Green Supermix (Bio-Rad). The data were analyzed using Bio-Rad CFX manager software. At3g18780 (Actin) and At4g05320 (UBQ10) were used as reference genes for normalization of gene expression levels in all samples, then the normalized gene expression levels were expressed relative to wildtype controls in each experiment.

**Table 2.**
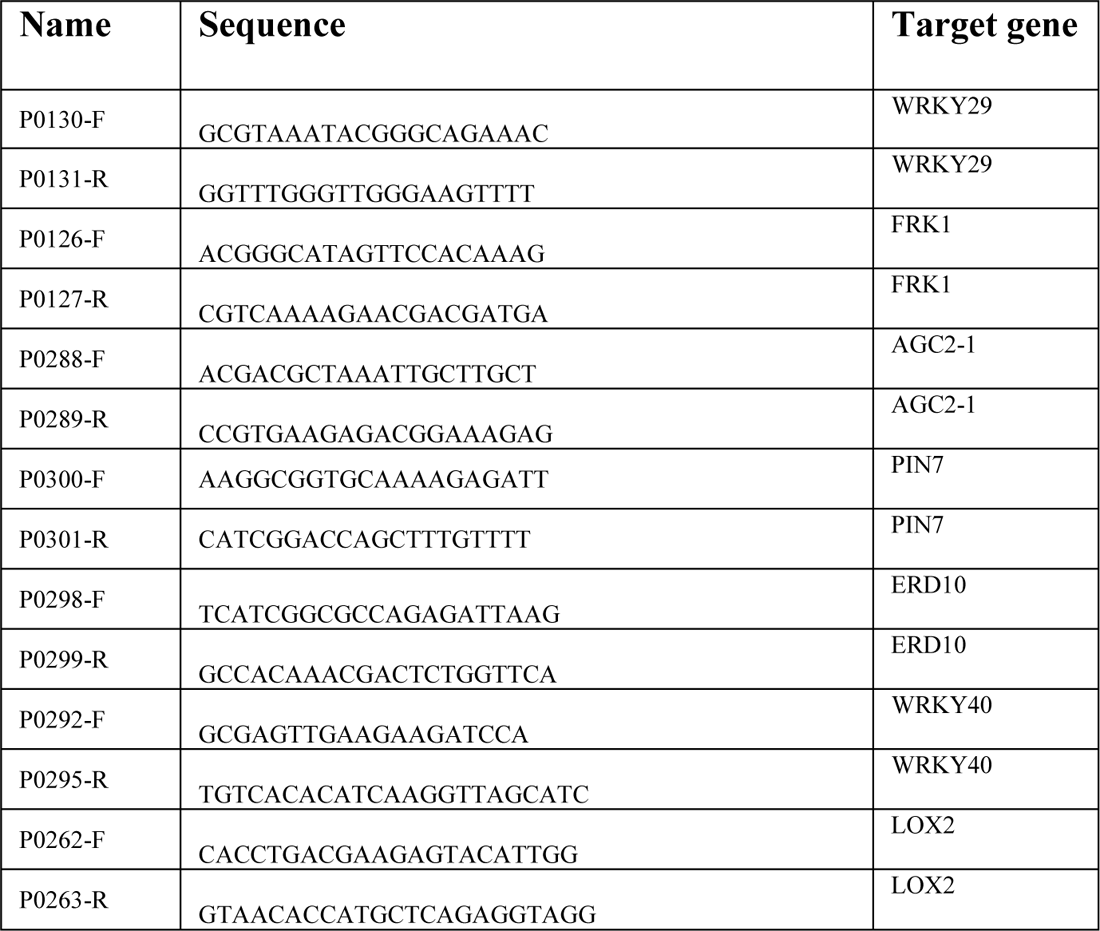
Primers Used For qPCR.

### Pathogen Assays

Plants were grown under short day conditions and spray-inoculated with *Pseudomonas syringae pv.tomato* DC3000) (*Pst*) at OD_600_= 0.2 with 0.04% (*v/v*) Silwet L-77. Disease symptoms were evaluated at 3 and 72 hours post-infection; leaf discs were sampled and bacteria were extracted. In total, three biological replicates were taken from 30 plants of each plant genotype. Bacteria were extracted with 10 mM MgCl_2_ containing 0.01% (v/v) Silwet L-77. The homogenates were plated on LB agar media containing rifampicin (50 mg/L) after a series of gradient dilutions, incubated at 28 °C for 48 hours, and bacterial colonies were counted. The infection level of the leaf samples from three biological replicates was reported as CFU/cm^2^.

*For Botrytis cinerea* infection, plants were grown short day conditions and inoculated by placing a 5 µL droplet of a suspension (5*106 spores/ml) on each rosette leaf. Disease symptoms were evaluated after 72 hours post infection, picture of the plants was taken and quantified using ImageJ Software. In total, three biological replicates of 9 plants each plant genotype by each taking 3 leaves per plant.

### ROS Burst Assays

ROS burst assays were carried out using the luminol-based luminescence method (Huang, Desclos-Theveniau et al., 2013). Briefly, 4 mm leaf discs of four weeks old *Arabidopsis* WT and mutant plants were incubated overnight (adaxial side up) in 150 µL of sterile water in 96-well plates (Thermo Fisher). The next day, water was replaced with 100 µL of reaction solution containing 50 µM of luminol (Sigma, St, Louis, Mo, USA), 10 µg/mL of horseradish peroxidase (Sigma, St, Louis, MO, USA), and supplemented with 1 µM of flg22 or water as a mock control. Luminescence was recorded at a 1 min intervals after the addition of flg22 for 40 min using a TECAN infinite 200 PRO microplate reader. The signal integration time was 0.5 sec and the ROS measurements were expressed as means of RLU (Relative Light Units).

### DAB and NBT stainings

Four-week-old fully developed *Arabidopsis* WT and mutant plants were stained for superoxide radical and hydrogen peroxide labeling, using nitroblue tetrazolium (NBT) (N6876, Sigma-Aldrich) or 3,3’diaminobenzidine (DAB) (D5637, Sigma-Aldrich) (Daudi et al., 2012), respectively. The leaves were then mounted on slides with 50% glycerol and photographs taken using a Nikon SMZ25 stereomicroscope.

### pTEpY Assays and Immunobloting

Total proteins were extracted from WT and *wlim2a Arabidopsis* seedlings upon cultivation on ½ MS plates for 14 days. The complete seedlings were crushed using a tissue lyser, resuspended in 200 µL SDS sample buffer, heated at 85° C for 10 minutes, centrifuged at 18,000 X g for 10 minutes at 4^0^ C, and the supernatant was collected. Using 10% SDS-polyacrylamide gels, total proteins were separated and electro-transferred onto PVDF membranes (BIO-RAD). The blots were blocked for one hour at room temperature with 5% (w/v) BSA in TBST before being incubated overnight at 4^0^ C with anti-phopho-p44/42 MAPK (diluted 1/75,000) in 2% (w/v) BSA in TBST (#4370, Cell Signalling). As a secondary antibody, HRP-conjugated goat anti-rabbit IgG was utilized (Promega). Goat antirabbit antibodies conjugated to horseradish peroxidase (HRP) were used as secondary antibodies. HRP activity was detected with a chemiluminescent reagent. Coomassie blue staining of blots was then carried out for protein visualization. Each immunoblotting analysis shown is representative of three independent biological repeats.

### RNA Sequencing Analysis

14-day-old WT and mutant *Arabidopsis* seedlings were grown in three independent biological replicates on ½ MS agar, moved to ½ MS liquid media overnight, and then treated for one hour with 1 µM flg22, a peptide synthesized by GeneScript, and water (mock). The sequencing was performed on HiSeq 4000 platform with a read length of 300 bp paired ends. Reads were quality controlled using FASTQC v0.11.5 (http://www.bioinformatics.babraham.ac.uk/projects/fastqc/). Trimmometric was used for quality trimming.

Parameters for read quality filter were set as follows: Minimum length of 36 bp; Mean Phred quality score greater than 30; Leading and trailing bases removal with the base quality below 3; sliding window of 4:15. TopHat v2.1.1 was used to alignment of short reads to the *Arabidopsis thaliana* TAIR 10, cufflinks for transcripts assembly, and differentially expression levels (fragment per kilobase of transcript per million mapped reads, FPKM). The differentially expressed genes (DEGs) were identified using Cufflinks and the limma package in R. Cuffdiff v2.2.1 (Trapnell et al., 2012) with quartile normalization to find the significant gene expression. Genes with 2-fold change and P-value <= 0.05 were considered as significantly different between samples with and without flg22 treatment. The hierarchical clustering of these genes was performed using Mev v4.8.1 (Howe, Sinha et al., 2011). Gene ontology (GO) classification was performed with the AgriGo (Du, Zhou et al., 2010) and DAVID Bioinformatic software (https://david.ncifcrf.gov/, Dennis et al., 2003). Venn diagrams are generated using (http://bioinfogp.cnb.csic.es/tools/venny/).

### Hormone Analysis

WT and mutant *Arabidopsis* seedlings were grown for two weeks on ½ MS agar, then moved to ½ MS liquid media overnight. The seedlings were then treated for an hour on the ½ MS with 1 µM flg22 peptide and water (mock). Phytohormone extraction was carried out as previously reported (Trapp et al, 2014). A Thermo Fisher TQS-Altis Triple Quadrupole Mass Spectrometer linked to a Thermo Scientific Vanquish MD HPLC system was used to quantify the chemicals using HPLC-ESI-SRM. The compounds were eluted using water (A) and acetonitrile (B) as mobile phases at 0.6 mL/min and in a gradient elution mode as follows: 10% B for 0.5 min, 10-55% B at 4.5 min, 55-100% B at 4.7 min, and 100% until 6.0 min. The temperature in the column was set to 55°C. To determine the statistical significance of three replicates, ANOVA was used, followed by the Tukey Test.

### Site-directed Mutagenesis

Wild type sequences of *WLIM2A* were amplified by PCR from *Arabidopsis* cDNA and cloned into PENTR-D-TOPO vector (**Table 3**). Mutated versions of *WLIM2A* were generated by site-directed mutagenesis using the Gene Art Site-Directed Mutagenesis System kit from ThermoFisher Scientific according to the manufacturer’s instructions reaction was carried. The primers were designed to introduce the mutations are listed in (**Table 3**). The introduction of mutations were confirmed using next generation sequencing (NGS) (**Table 4**) and the mutated transformed into final destination vector using the LR clonase (Life technologies). The constructs were transformed into *Agrobacterium* cells and using floral dip agrobacterium-mediated transformation construct were introduced into the homozygous *wlim2a-1* and *wlim2a-2* lines.

**Table 3.**
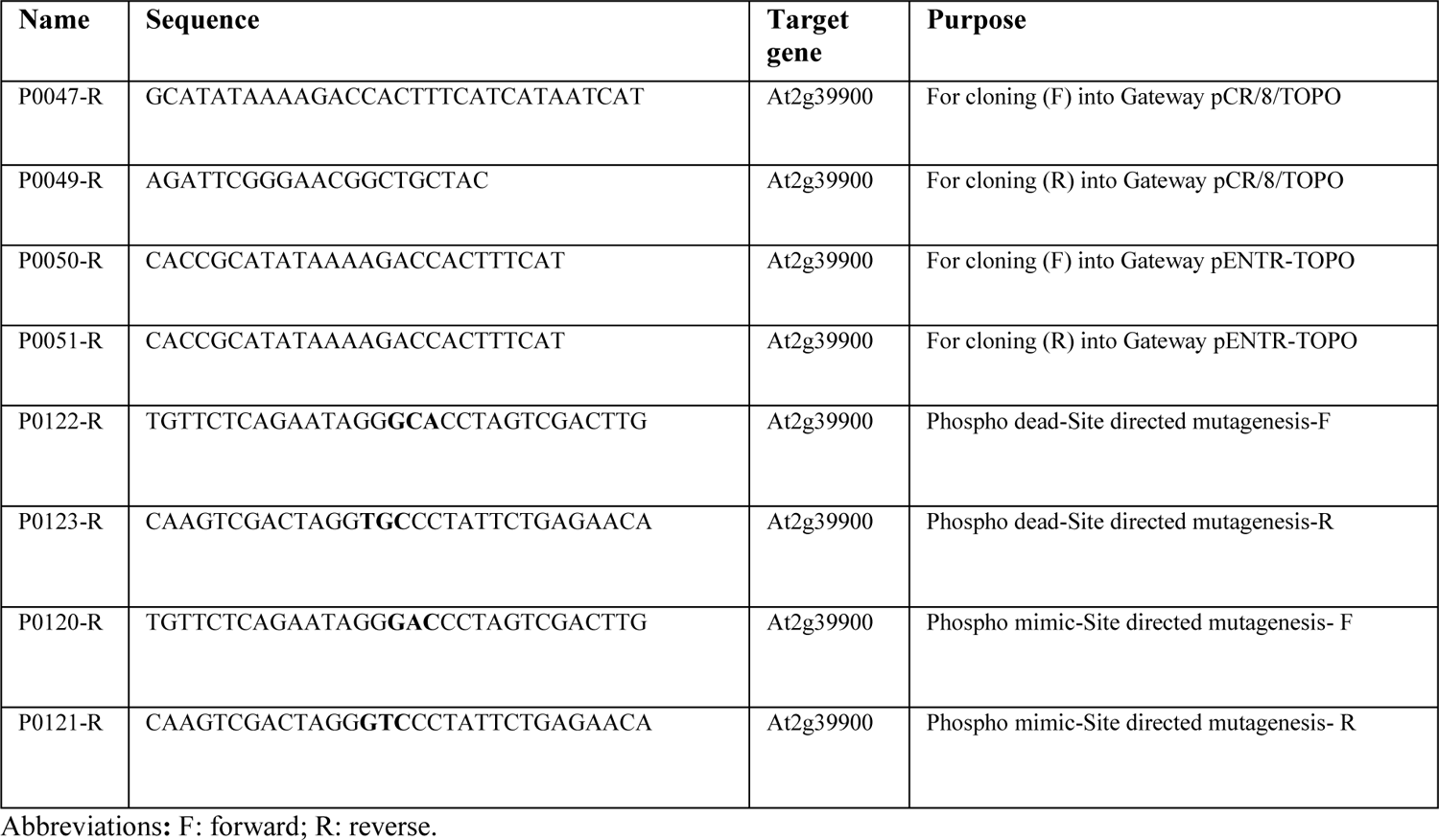
Primers Used For Cloning.

**Table 4.**
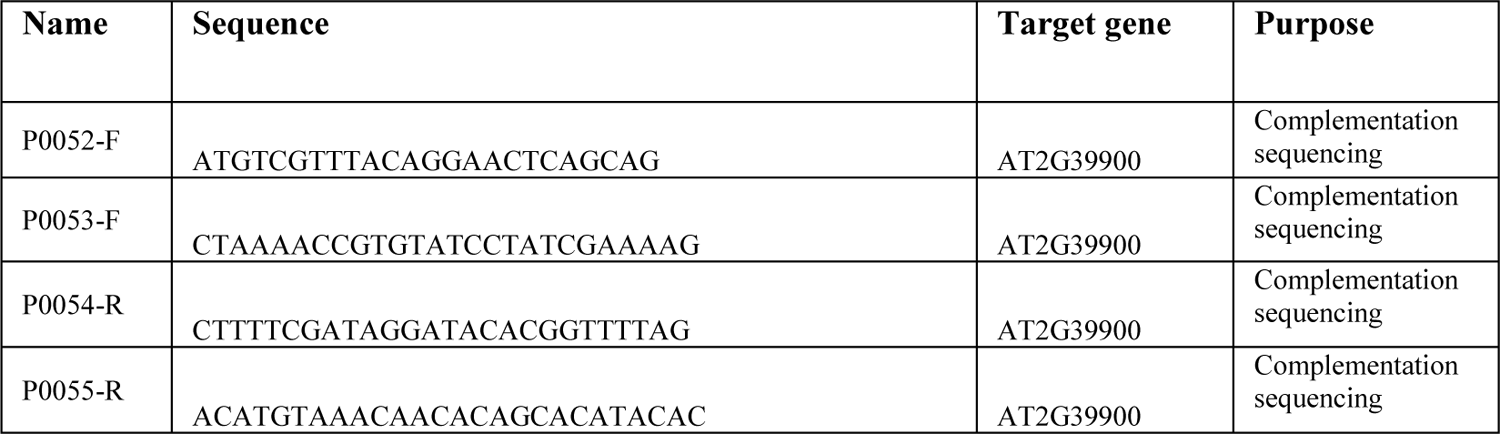
Primers Used For Sequencing.

### Bimolecular Fluorescence Complementation (BiFC)

To generate expression plasmids, coding sequences of the candidate genes, At1g11360, At2g39900 and MAPKs, were cloned into N or C-terminal part of YFPs under the promoter of 35S::GFP, in the pBIFC1, 2, 3 and 4 vectors. A range of positive and negative controls was prepared for the experiments. The construct carrying vector were transformed into *Agrobacterium tumefaciens* C58C1 strain. The cultures were grown on the LB medium with corresponding antibiotic selection marker for 24 h at 28°C. The culture was pellet down and resuspended with infiltration buffer (10 mM MgCl_2_, 10 mM MES pH 5.7, 150 µM acetosyringone) to an OD600 of 1.5 and kept in the dark for 3 h. The P19 viral suppressor of gene silencing was used to co-express with each combination to prevent silencing of transiently expressed proteins. Bacterial culture of each combination was mixed before infiltration. For a transient expression, the resultant bacterial suspension was directly infiltrated into *N. benthamiana* leaves. Three days after infiltration, the leaves were visualized using an upright LSM 710 Zeiss confocal microscope with a 20X objective (Plan-Apochromat, NA1.0) for GFP fluorescence. All images were obtained using the Argon laser with 514-nm excitation.

### Subcellular Localization Studies

The coding sequence of the candidate gene At2g39900, were cloned into p35S::C-GFP in the pGWB5 vector. The construct was transformed into *Agrobacterium tumefaciens* C58C1 strain. For a transient expression, the resultant bacterial suspension was directly infiltrated onto *N. benthamiana* leaves. An upright Zeiss LSM710 confocal microscope fitted with a 20X objective (Plan-Apochromat, NA1.0) was used to visualize the leaves with GFP fluorescence three days after infiltration. All images were acquired using the Argon laser at 514 nm.

### *In vitro* Kinase Assays

The purified recombinant proteins and constitutively active MAPKs were mixed together in kinase buffer. The kinase buffer contains (20 mM Tris-HCl pH 7.5, 10 mM MgCl_2_, 5 mM EGTA, 1 mM DTT, and 50 µM ATP) and the reaction mixture is incubated at ambient temperature for 30 min. To stop the reaction, SDS-sample buffer was added and followed by incubation at 95°C for 10 min. The protein samples were resolved using SDS-PAGE. SimplyBlueTM SafeStain (Novex cat. no. LC6065) was used to stain the gel, and the band corresponding to the protein of interest was excised out, cut into small pieces of 0.5 mm^3^, and destained with four successive washes of 15 minutes each with ACN and 100 mm NH4HCO3. Proteins were reduced at 37 °C for 1 hour with 10 mm Tris(2-carboxyethyl)phosphine (TCEP, C-4706 Sigma) in 100 mm NH_4_HCO3, followed by alkylation at ambient temperature for 30 minutes with 20 mm S-Methyl methanethiosulfonate (MMTS, 64306 Sigma). Proteins were then digested overnight at 37°C with trypsin (Porcine trypsin, Promega, Fitchburg, WI). After stopping the digestion with 1% formic acid, the peptides were recovered by incubating the gel pieces in ACN. The desalted peptide solution was analyzed by LC-MS/MS using a C18 ZipTip® (Millipore, Burlington, MA, Cat. No. ZTC18S096).

## Results

### Validation of *in vivo* Phosphorylation Sites of the MAPK Substrate WLIM2A

WLIM2A (At2g39900) was identified as a putative MAPK target that is differentially phosphorylated after treatment with flg22, a microbe-associated molecular pattern (MAMP). Comparative phosphoproteomics of WT and *mapk* (*mpk3*, *mpk4,* and *mpk6*) mutants suggested a possible involvement of WLIM2A in PAMP-triggered immunity (**Fig. S3;** Rayapuram et al., 2018). In order to further investigate the role of WLIM2A in MAPK signaling, we first investigated whether WLIM2A can directly interact with any of the three MAPKs MPK3, MPK4, and MPK6. For this purpose, we tested the interaction between WLIM2A and the three immune MAPKs (MPK3, MPK4, and MPK6) *in vivo* using bimolecular fluorescence complementation (BiFC) assays. The proteins were transiently co-expressed in leaf epidermal cells of *Nicotiana benthamiana.* Confocal images confirmed that WLIM2A strongly interacts with MPK3 in both the nucleus and cytoplasm, while MPK4 and MPK6 interaction was less observed *in planta* (**Fig. 1A**). Moreover, YFP fluorescence formed by the interaction of WLIM2A with MPK3, MPK4, and MPK6 was predominantly observed in the cytoplasm.

**Figure 1.**
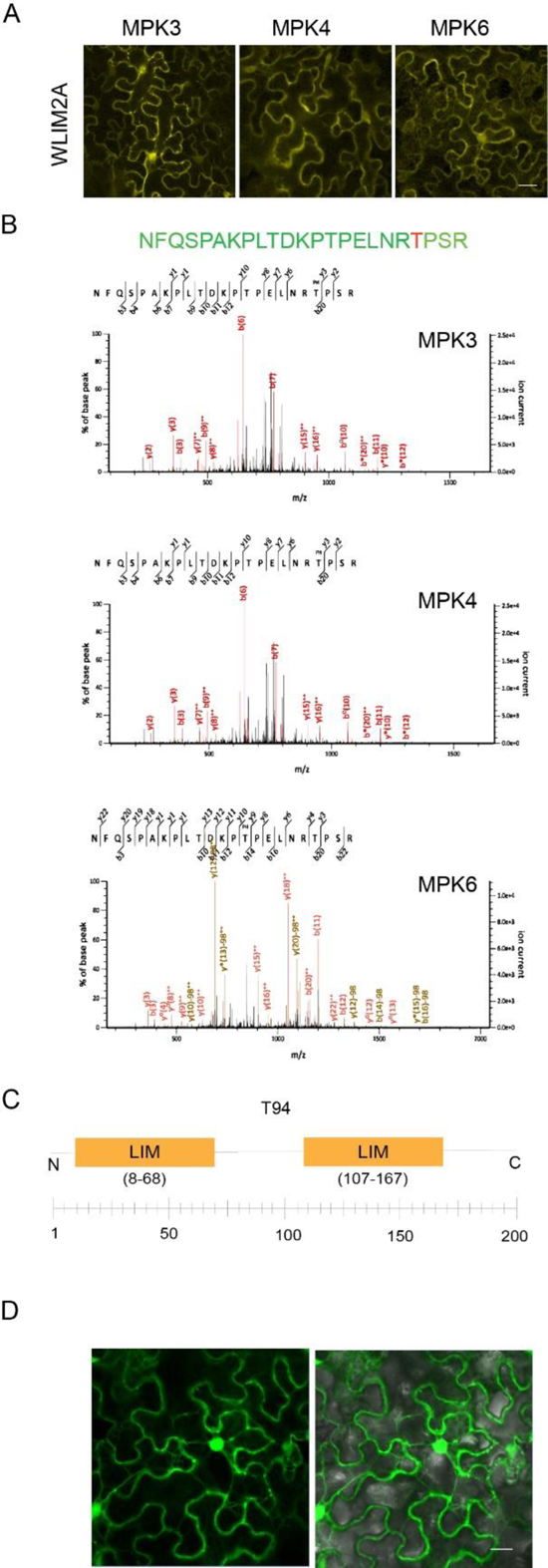
(**A**) BiFC investigation using MPK3, MPK4, and MPK6 in *N. benthamiana* leaf epidermal cells of the MAPK candidate substrate WLIM2A. Laser scanning confocal microscopy was used to detect YFP fluorescence. The scale bar measures 20 µM. (**B**) Analysis of samples from an *in vitro* kinase test with MPK3, MPK4, and MPK6 kinase and WLIM2A fragment by mass spectrometry. Two phosphorylation sites were discovered in the spectrum: 1TP (indicated in red), and 1SP (marked in green). (**C**) WLIM2A protein structure shows the relative position of two lim domains and position of the phosphosite obtained from the phosphoproteomics screen. (**D**) Subcellular localization of WLIM2A tran-siently expressed MAPK candidate substrate coupled to GFP in *N. benthamiana* cells driven by the 35S promoter. The tagged proteins were expressed in 4-week-old tobacco plants, and the localisation was observed using laser scanning confocal microscopy between 48 and 72 hours after infiltration. The scale bar measures 50 µM.

To further characterize WLIM2A, we tested the capacity of the MAPKs to phosphorylate WLIM2A at the identified T94 *in vivo* phosphorylation site (Rayapuram et al., 2018) (Fig. 1B **&** Fig. 1C). We performed *in vitro* kinase assays with recombinant WLIM2A and constitutively active MPK3, MPK4, and MPK6 proteins. LC-MS/MS revealed that WLIM2A was phosphorylated at T94 by all three MAPKs *in vivo*, confirming that WLIM2A is a direct MAPK target (Fig. 1B). We next examined the subcellular localization of GFP-tagged WLIM2A protein expressed in *N. benthamiana* leaf epidermal cells. Confocal microscopy images showed that WLIM2A localized to both the cytoplasm and nucleus (Fig. 1D). These data align with our previous phosphoproteomic study (Rayapuram et al., 2018), which identified WLIM2A as a potential MAPK target.

### Targeted Mutagenesis of *WLIM2A* using the CRISPR-Cas9 System

In order to characterize the function of WLIM2A, we generated mutants (knockout) using CRISPR-Cas9 technology. Gene-specific primers were designed to amplify the *WLIM2A* fragment from WT Col-0 and CRISPR-Cas9 *wlim2a* lines. We screened for mutations in the coding region of the gene by sequencing. We obtained lines that had either an addition of guanine or thymine in the coding region that resulted in a frameshift mutation and premature termination of the WLIM2A protein (Fig. 2A**, S4A &B**). To avoid off-target mutations, we selected subsequent generations for the lack of Cas9 by PCR-based genotyping using gene-specific primers (**Fig. S4C**). Both *wlim2a-1* and *wlim2a-2* exhibited normal developmental phenotypes and no obvious morphological or developmental defects compared to WT plants (Fig. 2B). Determination of the relative transcript levels by qPCR indicated that both loss-of-function transcripts are present at low levels in *wlim2a-1* and *wlim2a-2* mutants (Fig. 2C).

**Figure 2.**
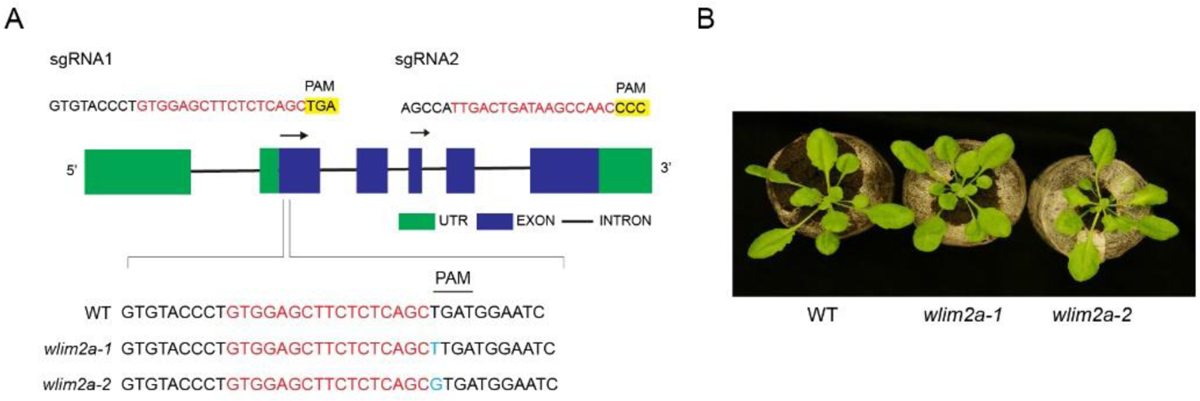
(**A**) This figure shows CRISPR-Cas9 genome-editing strategy in *Arabidopsis thaliana*, based on two sgRNA targeting *WLIM2A* and sequencing reads from selected mutant alleles from T0 transformants show an addition of thymine or guanine. (**B**) WT (Col-0), *wlim2a-1*, and *wlim2a-2* mutants, have the same morphological phenotype. Plants cultivated in jiffy peat pellets for four weeks are displayed.

### Immune responses to infection by a virulent bacterial pathogen is compromised in *WLIM2A* **plants**

We investigated the role of WLIM2A in plant immunity upon challenge with *Pst.* We spray-inoculated *wlim2a-1* and *wlim2a-2* plants with the pathogenic strain *Pst*. We observed enhanced susceptibility of both of the *wlim2a* mutant lines compared to WT at 72 h post infection (Fig. 3A). We measured the levels of salicylic acid (SA), a key hormone that regulates plant immunity to bacterial pathogens. The data showed significantly lower SA levels in the *wlim2a-1* mutant compared to WT in both untreated and flg22 treated plants (Fig. 3B). Therefore, this observation shows that WLIM2A plays a positive role in regulating disease resistance to bacterial pathogen.

**Figure 3.**
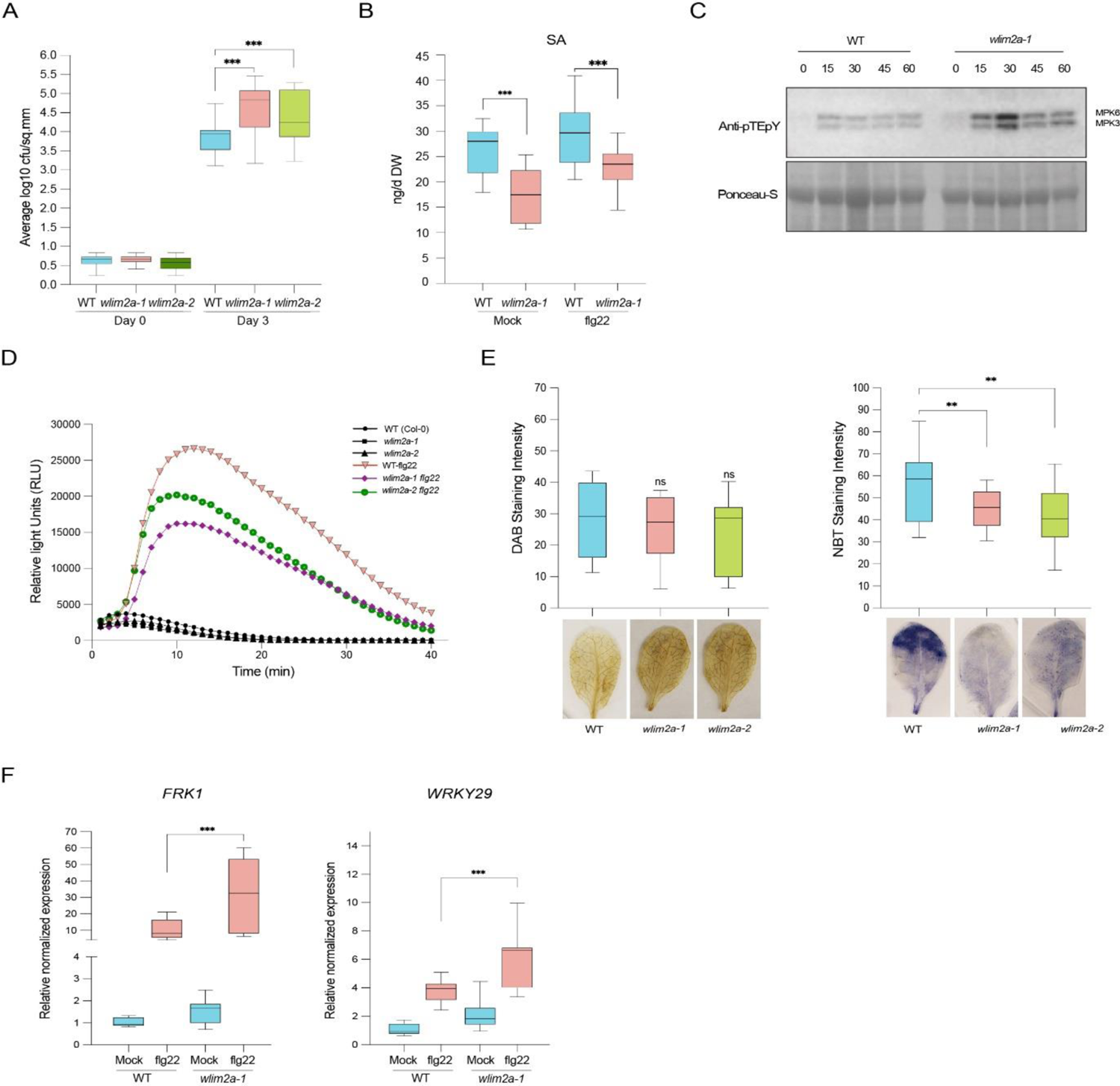
(**A**) *Arabidopsis* plants infected with *Pseudomonas syringae pv. tomato* DC3000 (Pst) in *wlim2a-1, wlim2a-2*, and WT (Col-0). Four-weeks-old seedlings were spray-inoculated with Pst or mock. Pst colonies were counted 3 and 72 hours after inoculation (hpi). The vertical bars represent the standard error for the three biological replicates with 16 plants each. Student’s t test, p<0.05 was used to determine statistical significance. The median is shown by the lines in the boxes. (**B**) SA levels were quantified by LC-MS/MS in Col-0, *wlim2a-1*, and *wlim2a-2* lines after treatment with or without flg22 for 1-hour. 18 plants for each line were used in three biological replicates with 3 seedlings at 100 mg/tube each. (**C**) MPK3 and MPK6 are rapidly and transiently activated by flg22 in mock and *wlim2a-1* mutants. Col-0 and the *wlim2a-1* mutant were incubated in water overnight before being exposed to 1 µM flg22 for the specified amount of time. By utilizing an anti Tp-E-Yp antibody against the phosphorylated version of ERK2, western blotting was used to detect the phosphorylation of MPK3 and MPK6. Equal loading of proteins was shown by Coomassie staining. (**D**) In *wlim2a* mutants, flg22-induced ROS generation is reduced. Leaf disks from four-week-old *Arabidopsis thaliana*, Col-0, *wlim2a-1*, and *wlim2a-2* plants were collected and treated with 1 mM flg22 or mock. Over 40 minutes, a luminol-based assay detected ROS accumulation as Relative Light Units (RLU). Three independent experiments generated similar findings (n=12/treatment). (**E**) Evaluation of H2O2 levels in Col-0 and mutants using 3,3’-diaminobenzidine staining (DAB) and O2^-^ levels in mutants using NitroBlue Tetrozolium (NBT) staining in comparison to WT Col-0. (**F**) PTI marker gene relative expression levels in WT (Col-0)*, wlim2a-1,* and *wlim2a-2*. UBIQUITIN and ACTIN were used as reference genes to normalize the data. The X-axis displays the expression of *FRK1* and *WRKY29* in mock- and flg22-treated *Arabidopsis* lines. The Y-axis displays the changes in gene expression levels shown on a log2 scale. The error bars show the standard error (SE), and an asterisk (*) denotes a p-value of less than or equal to 0.1, 0.05, or 0.001 for the Student’s t-test. Three independent experiments, each with a n = 9. The standard error (SE) of the data is shown by the error bars (* denotes P<0.05, *** P>0.001, Student’s t-test).

As both *wlim2a* mutant alleles exhibited increased susceptibility to *Pst*, we examined whether the cellular responses involved in PTI were affected in the mutant lines. Increased production of reactive oxygen species (ROS) is one of the earliest immune responses following PAMP perception. Another hallmark of PTI is the activation of various MAPKs, including MPK3, MPK4 and MPK6. Surprisingly, *wlim2a-1* mutant seedlings showed higher MPK3, 4 and 6 activation levels than WT after 15 and 30 mins of flg22 elicitation (Fig. 3C). However, flg22-induced ROS production was lower in both *wlim2a-1* and *wlim2a-2* compared to WT plants (Fig. 3D).

To study the role of WLIM2A in stress responses, WT plants and *wlim2a-1* and *wlim2a-2* mutants were stained for hydrogen peroxide using diamino-benzidine 3,3’-diamino-benzidine (DAB) and for superoxide radicals using nitroblue tetrazolium (NBT). Polymerization of DAB and NBT can be spotted as a brown precipitate in the presence of hydrogen peroxide and as a blue precipitate in the presence of superoxide, respectively. Both *wlim2a-1* and *wlim2a-2* mutants accumulated similar H_2_O_2_ (DAB staining) but less O ^-^ (NBT staining) compared with WT plants (Fig. 3E). We then examined the role of WLIM2A in PAMP-triggered transcriptional responses of known PTI marker genes. Quantitative RT-PCR (qRT-PCR) analysis showed that the basal transcript levels of *FRK1* and *WRK29* were higher in *wlim2a-1* in both untreated and flg22-treated plants when compared with WT plants (Fig. 3F).

### Global transcriptomic profile shows that WLIM2A regulates defense gene expression

To characterize the function of WLIM2A in the regulation of gene expression, we carried out a global transcriptome analysis of *wlim2a-1* mutant plants in response to flg22 treatment. The expression levels were normalized using the fragment per kilobase of million mapped reads (FPKM). Differentially expressed genes (DEGs) were identified based on a minimum of a two-fold change in gene expression and false discovery rate (FDR) value (P ≤ 0.05). We obtained 1816 DEGs in response to flg22, 148 upregulated genes, and 931 downregulated genes in the WT, whereas in *wlim2a-1* we obtained 554 upregulated and 778 downregulated genes in response to flg22 (Fig. 4A**; S5**).

**Figure 4.**
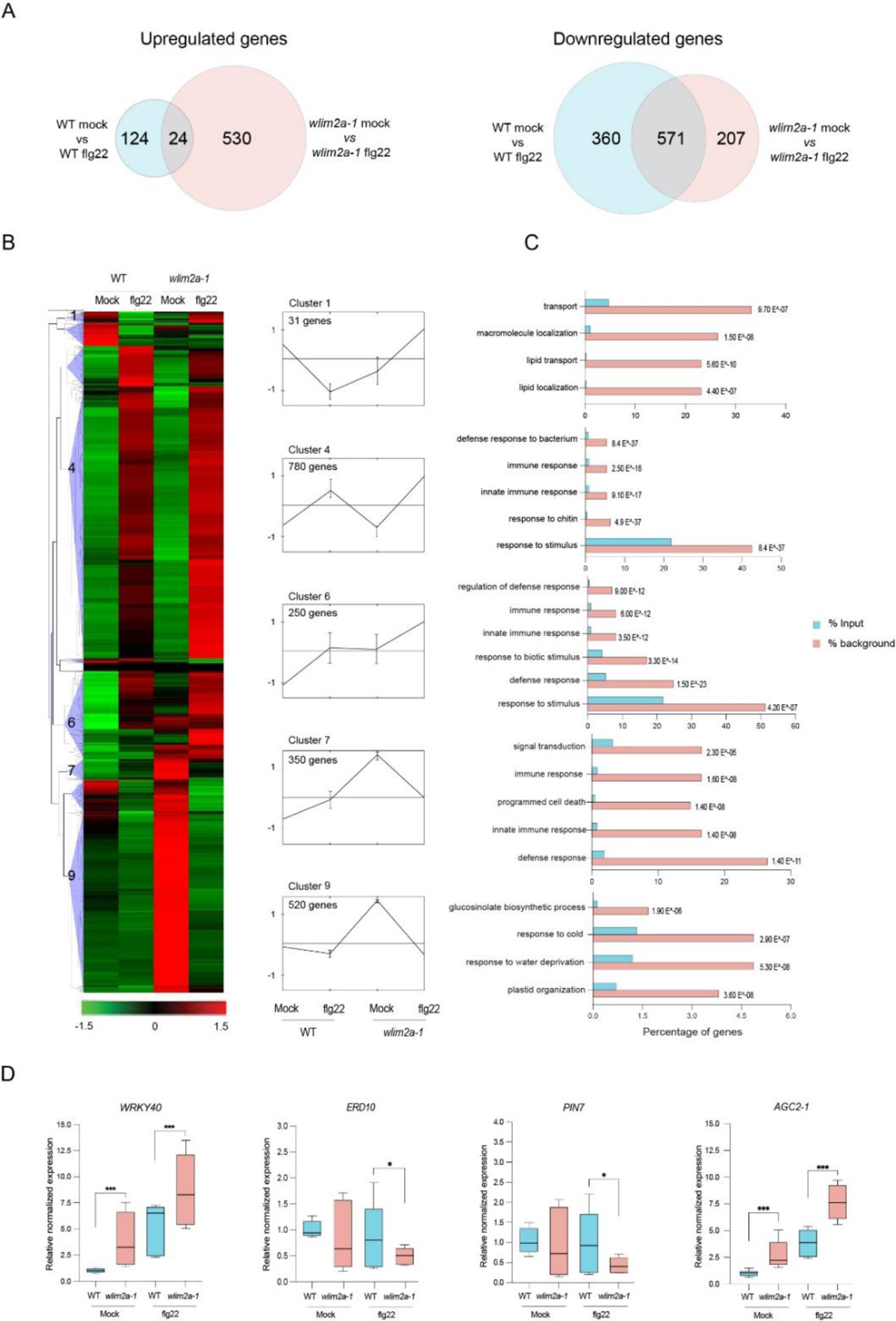
(**A**) Venn diagram displaying the overlap of differentially expressed genes (DEGs) identified by RNA-Seq between WT mock vs WT-flg22 and *wlim2a-1* mock vs *wlim2a-1* flg22. (**B**) The DEGs in WT mock, WT-flg22, *wlim2a-1* mock, and *wlim2a-1* flg22 were clustered hierarchically into ten groups. The values are provided as log 2-fold changes (P<0.01). In the clustering, the average linkage method and Pearson correlation (MEV4.0) were applied. The red and green colours represent up- and down-regulated expression, respectively. The black lines reflect the average relative expression levels, whereas the gray lines represent the relative expression of each gene in clusters 1,4,6,7, and 9. (**C**) Functional Enrichment of DEGs between *Arabidopsis* Col-0 (WT) mock, *wlim2a-1* mock, WT flg22, and *wlim2a-1* flg22 in the GO databases. Blue represents input, and orange represents the reference control. RNA-seq expression data validation by (RT-qPCR). Actin and ubiquitin were used to standardize the signal intensity for each transcript. (**D**) Gene expression in WT mock, WT flg22 treated, *wlim2a-1* mock, and *wlim2a-1* flg22 treated is shown on the X-axis. The Y-axis displays the log2 scaled variations in gene expression levels. The error bars represent (SE, standard error, one asterisk (*) indicates P<0.05, three (***) P>0.001, Student’s t-test).

To gain a better understanding of the genes affected by deletion of WLIM2A, we performed hierarchical clustering analysis of flg22-induced genes followed by GO enrichment analysis. In total, 1816 DEGs were up and downregulated were displayed between WT and *wlim2a-1* treated samples with P ≤ 0.05. The clustering analysis of all DEGs was carried out using the Euclidean distance method associated with complete linkage. Ten clusters were identified based on similar expression patterns (Fig. 4B). Gene Ontology (GO) functional enrichment of DEGs was carried out for the different clusters using AgriGo analysis.

Focusing on three specific clusters of interest, cluster 1 comprised 31 DEGs that were highly upregulated in *wlim2a-1* upon flg22 treatment but not in WT. Enrichment analysis of the GO categories of cluster 1 revealed genes associated with response to localization and macromolecule transport. Cluster 7 and 9 genes were upregulated in untreated *wlim2a-1* plants. Cluster 7 contains 350 DEGs involved in signal transduction, defense response, and cell death, whereas cluster 9 contains 520 DEGs that are related to the glucosinolate process, response to water and cold, and plastid organization (Fig. 4C).

In order to validate the RNA-seq analysis, we performed qRT-PCR on several of the DEGs. We observed differential upregulation of transcriptional factors *WRKY40* and *AGC2-1* before and after flg22 treatment in *wlim2a-1* compared to WT plants (Fig. 4D). The expression of *ERD10* and *PIN7* was repressed after flg22 treatment in *wlim2a-1* compared to WT plants (Fig. 4D).

### Loss of WLIM2A function enhances basal defense against *B. cinerea*

The transcriptome data encoded many components of pathways involved in the production or signaling of phytohormones. So, we assessed the concentrations of salicylic acid (SA) and jasmonic acid (JA), two important hormones that control plant immunity. We found no changes of JA, but increased levels of the active jasmonate, JA-Ile, in *wlim2a-1* the mutant relative to WT (Fig. 5A). To determine the role of the WLIM2A in fungal defense, we examined the response of WT, *wlim2a-1* and *wlim2a-2* mutant plants to *B. cinerea*. = *wlim2a-1* mutant plants were more resistant than WT (Fig. 5B). Higher expression levels of the JA and *B. cinerea* related genes (such as *BIK1, and LOX2*) were observed in *wlim2a-1* mutants when compared to WT in flg22 treated plants (Fig. 5C). These findings support that WLIM2A plays a negative role in *B. cinerea* defense.

**Figure 5.**
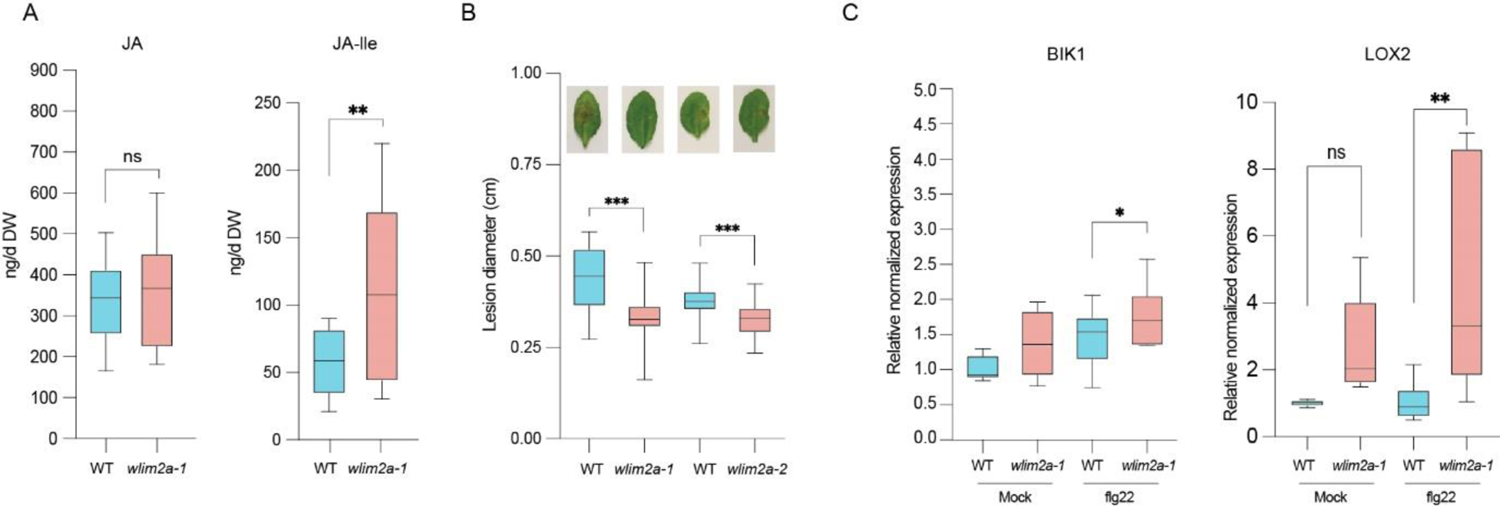
(A) Quantitative study of JA and JA-lle levels in Col-0, *wlim2a-1*, and *wlim2a-2* lines using LC MS/MS with or without flg22 treatment for 1 hour. Plants of three seedlings of 100mg/tube and 18 plants for each line three biological replicates. (B) *Arabidopsis WLIM2A* mutations enhanced the plant more resistance to the fungus *B. cinerea*. *A. thaliana* plants four weeks old were injected with 5 L droplets of fungal spores (5 ×10^5^ spores mL^−1^). The area of lesion in the leaves of Col-0 and *wlim2a* mutant plants was assessed 48 hours later. ImageJ software was used to determine the size of the lesions. Vertical bars represent the standard error for three biologically independent experiments (n=8 for each). 3 leaves per plant, with an average of 8 plants per replicate. (C) RNA-seq expression data validation (RT-qPCR). The intensity of each transcripts signal was normalized using actin and ubiquitin. The X-axis shows gene expression in WT mock, WT flg22 treated, *wlim2a-1*, mock, and *wlim2a-1*, flg22 treated *Arabidopsis*. The error bars represent (SE, standard error, one asterisk (*) indicates P<0.05, three (***) indicates P>0.001, Student’s t test).

### Essential role of WLIM2A phosphorylation in immunity

To study the function of WLIM2A protein by MAPK phosphorylation at the identified T94 phosphorylation site, we generated stable *WLIM2A-WT* (ProUbi::*WLIM2Awt-GFP), WLIM2A-PM* phosphomimic (ProUbi::*WLIM2AT94D-GFP)* and *WLIM2A-PD* phosphodead (ProUbi::*WLIM2AT94A-GFP)* lines in the *wlim2a-1* mutant background by site directed mutagenesis. Phenotypically, we found no differences to WT in growth or development of the *WLIM2A-PD or WLIM2A-PM* lines (Fig. 6A).

**Figure 6.**
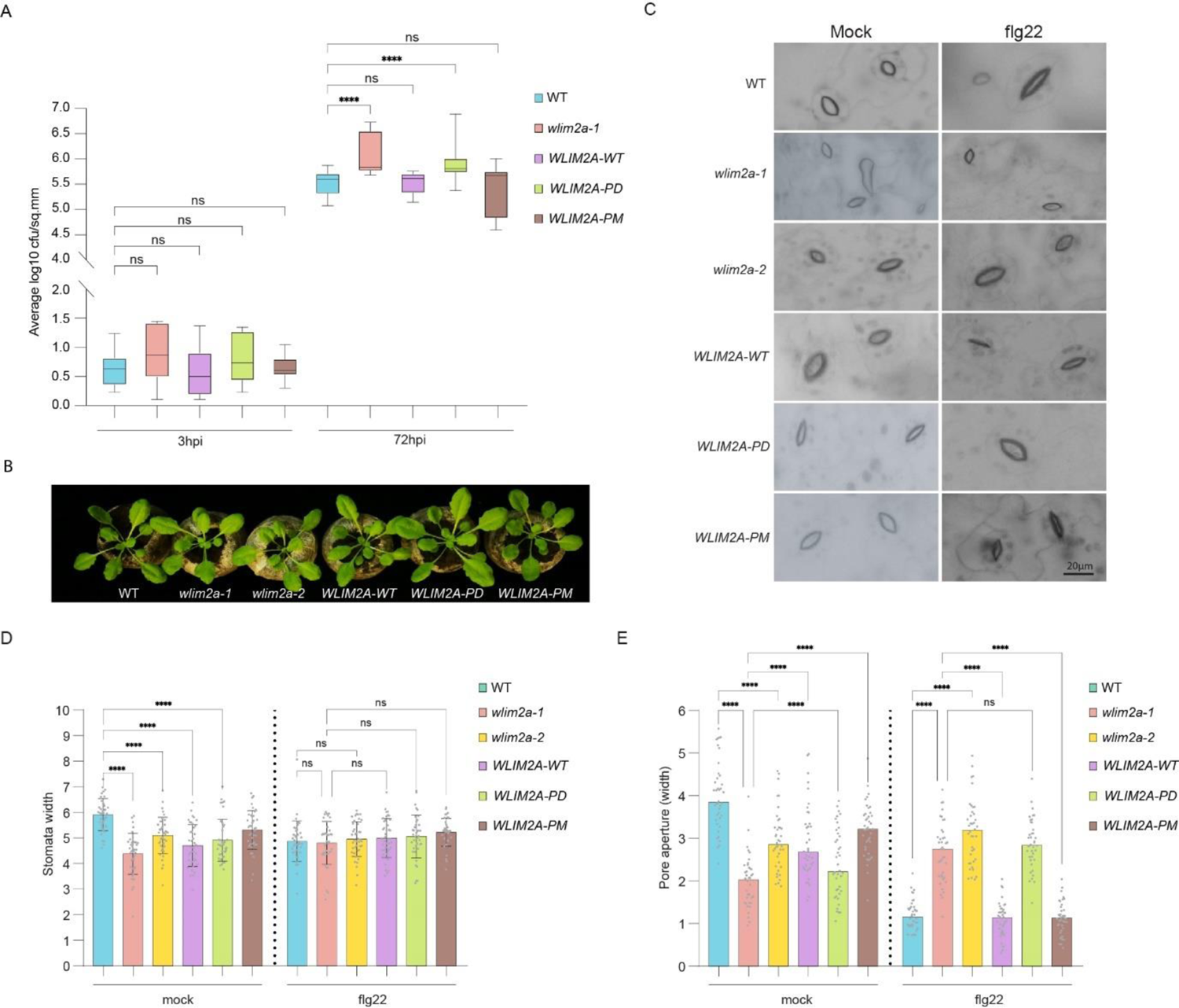
WLIM2A plays an essential role in stomatal immunity in *Arabidopsis*. (**A**) Four-weeks-old Arabidopsis seedlings of *wlim2a-1, wlim2a-2*, *WLIM2A*-*WT*, *WLIM2A*-*PD, WLIM2A*-*PM* and WT (Col-0) were spray-inoculated with *Pseudomonas syringae pv. tomato* DC3000 bacterial suspension (conc) or mock. Susceptible to the bacterial pathogen *Pseudomonas syringae pv. tomato* DC3000 of *wlim2a-1*, *wlim2a-2* mutants in comparison to Col-0. *WLIM2A*-*WT* showed recovery of the pathogen phenotype. The colonies were counted 3 and 72 hours after inoculation (hpi). The vertical bars represent the standard error for the three biological replicates with 16 plants each. (**B**) Representative images of the stomatal closure in the MAMP-induced stomatal defense in WT, *wlim2a-1*, and *wlim2a-2*, *WLIM2A*-*WT*, *WLIM2A*-*PD, and WLIM2A*-*PM*. (**C&D**) Stomatal aperture measurement of the WT, *wlim2a-1*, and *wlim2a-2*, *WLIM2A*-*WT*, *WLIM2A*-PD, and *WLIM2A*-*PM*. Scale bars measures 20 µM. Sample sizes in (b, c) are indicated. The error bars represent (SE, standard error, one asterisk (*) indicates P<0.05, three (***) indicates P>0.001, as determined by one-way ANOVA with Tukey’s multiple comparisons test.

We then investigated the role of WLIM2A phosphorylation with respect to defense against the pathogenic bacterial strain *Pst.* We spray-inoculated the different lines with *Pst* and compared infection levels to WT at 72 h post infection. In contrast to the enhanced infection levels in *wlim2a-1*, we observed similar susceptibility in *WLIM2A-WT* and *WLIM2A-PM* as in Col-0 (Fig. 6B). In contrast, *WLIM2A-PD* had similar *Pst* infection levels as the *wlim2a-1* mutant (Fig. 6B). First, these data confirm that the expression of *WLIM2A-WT* complements the *wlim2a-1* mutant phenotype. Second, they show that MAPK directed phosphorylation of WLIM2A plays a key role in pathogen resistance.

### Role of WLIM2A in Stomatal Immunity

Upon pathogen attack, stomates rapidly close as a primary defense mechanism to inhibit pathogen entrance into the leaf apoplast. We observed at 3 hpi increased bacterial growth on *wlim2a-1* and *WLIM2A-PD* mutants, which was not seen in WT or *WLIM2A-WT* and *WLIM2A-PM* lines. Although these differences in bacterial titers were not statistically significant, this trend prompted us to investigate a possible role of WLIM2 in stomatal physiology and stomatal defense (Fig. 6 **A**). For this purpose, we treated plants with flg22, which triggers stomatal closure, or with H_2_O as mock control. Under mock treatment, we observed differences in stomatal opening of WT, *wlim2a-1, wlim2a-2, WLIM2A-WT*, *WLIM2A-PD,* and *WLIM2A-PM*. Upon flg22 treatment, stomata of WT, *WLIM2A-WT* and *WLIM2A-PM* responded by closure, but not in the *wlim2a-1 and WLIM2A-PD* lines (Fig. 6 **B-D**). Together, these findings indicate that WLIM2A is crucial for the stomatal immunity response in *Arabidopsis*.

## DISCUSSION

Plant MAPKs are activated and play an important role in response to several biotic and abiotic stresses (Nakagami et al., 2005), but our knowledge of the respective MAPK substrates is limited. MAPK-dependent phosphorylation of their substrates and their subsequent functional characterization is vital to the understanding of plant stress responses. Therefore, in this study, we explored the functions of the putative MAPK target WLIM2A.

Here, we report that WLIM2A functions as a regulator of immunity in *Arabidopsis*. Notably, our phenotypic studies of *wlim2a* mutants showed that WLIM2A negatively regulates disease resistance against *Pst*. WLIM2A is known to regulate gene transcription in the nucleus (Moes et al., 2013). We analyzed the transcriptional changes following flg22 treatment using RNA-seq. We found that genes encoding WRKY40, a regulator of SA synthesis, NPR3, a receptor of SA immune signaling, and CBP60G, another regulator of SA, were upregulated in the *wlim2a* mutant after flg22 treatment. In addition, increased susceptibility to *Pst* DC3000 was observed in the *wlim2a* mutant. These results suggest that SA is regulated by WLIM2A and is involved in the regulation of the defense response.

Defense-related genes, including *WRKY27*, *PEN2*, *WRKY40*, *MYB51*, and *ERF6*, were upregulated in the flg22-treated WT and *wlim2a-1* mutant when compared to mock treated. MPK3 and MPK6 have an essential function in the induction of camalexin, a major phytoalexin in *Arabidopsis*. *Botrytis cinerea* infection activates MPK3/MPK6, promotes indole-3yl-methylglucosinolate (I3G) biosynthesis which is converted into 4 methoxindole-3-yl-nethylglucosinolate (4MI3G). MPK3 and MPK6 regulate the function of MYB51, a key regulator of IGS biosynthesis, through ETHYLENE RESPONSE FACTOR6 (ERF6), which is a substrate of MPK3/MPK6 and a target of PENETRATION2 (PEN2)-dependent chemical defense in plant immunity (Meng X et al., 2013; Juan Xu et al., 2016). MPK3/MPK6 phosphorylate ERF6 and thereby promote the biosynthesis of indole glucosinolates (Juan Xu et al., 2016). This result supports the idea that WLIM2A might control glucosinolate biosynthesis and is involved in plant immunity, although further validation and studies are required. The expression of *PIN7* and *LOX2* was significantly increased in the *Arabidopsis wlim2a* mutant, whereas these genes were downregulated after flg22 treatment in the *wlim2a* mutant, and the WT. JA and auxin may interact positively in plant resistance to the necrotrophic pathogen *B. cinerea* (Kazan et al., 2009). PIN7 is an auxin transporter and LOX2 is involved in JA biosynthesis. JA-dependent defense signaling could be part of the auxin-mediated defense mechanism involved in resistance to necrotrophic pathogens. These results suggest that the altered susceptibility phenotype of *wlim2a* mutants to infection by bacterial and fungal pathogens involves the transcriptional regulation of defense-related genes.

Four families of actin binding proteins (ABP) are involved in the formation or maintenance of the actin bundles in plant cells: fimbrin, formin, villin, and two LIM domain-containing proteins (LIM) (**Fig. S7**). Little is known about the function of LIM proteins. During plant innate immunity, actin cytoskeleton remodeling may require several fundamental cellular responses, including transcriptional activation, generation of ROS and antimicrobial compounds, directed vesicle trafficking, and fortification of the cell wall (Day et al., 2011). Treatment with the MAMP elf26 showed a significant increase in filament abundance and altered filament dynamics. The disruption of the actin cytoskeleton leads to increased plant pathogen resistance in a SA-dependent manner (Matouskova et al., 2014; Leontovycova et al., 2019).

Our quantitative large-scale phosphoproteomics studies revealed that WLIM2A is phosphorylated *in vivo* at T94 by MPK3, MPK4, and MPK6 (Rayapuram et al., 2017). Besides acting as a nuclear transcription factor, Hoffmann et al. (2014) previously showed that LIM can also crosslink cytoplasmic actin filaments. Therefore, protein phosphorylation may be involved in the subcellular shuttling of LIM from the cytoplasm to the nucleus, and thereby activate gene transcription in the nucleus (Srivasta et al., 2015). Our pathogen assays established that WLIM2A is a regulator of stomatal immunity and that MAPK phosphorylation of WLIM2A is the underlying mechanism to regulate stomatal closure. MPK3, MPK4, and MPK6 have been shown to be highly expressed in guard cells (Meng et al., 2013). Mutants of WLIM2A, *wlim2a-1*, and *WLIM2A-PD* fail to close stomata in response to pathogen attack, WT, *WLIM2A-WT* and constitutively phosphorylated *WLIM2A-PM* close stomata (Fig. 6C). Consistent with earlier research on actin binding and bundle actin filaments (Eun et al., 1997; Zou et al., 2019), actin severing, and failure of actin rearrangement will compromise stomatal closure (Kumar et al., 2004). In contrast, AtLIMs are known to stabilize the actin filament bundling (Thomas et al., 2007). During the pathogen infection, actin array undergoes reorganization to regulate the stomata opening or closure (Qian et al., 2019; Zheng et al., 2019), and molecular mechanism of the actin dynamics in the stomata requires further investigation. In our studies, we demonstrated that WLIM2A is rapidly phosphorylated upon flg22-activation of MAPKs. Combined genetic and biochemical evidence suggests that MAPK-induced phosphorylation of WLIM2A initiates stomatal immunity. Future research should establish the role of WLIM2A phosphorylation in actin dynamics and whether this is causal in regulating PAMP-induced stomatal closure and defense (Fig. 7).

**Figure 7.**
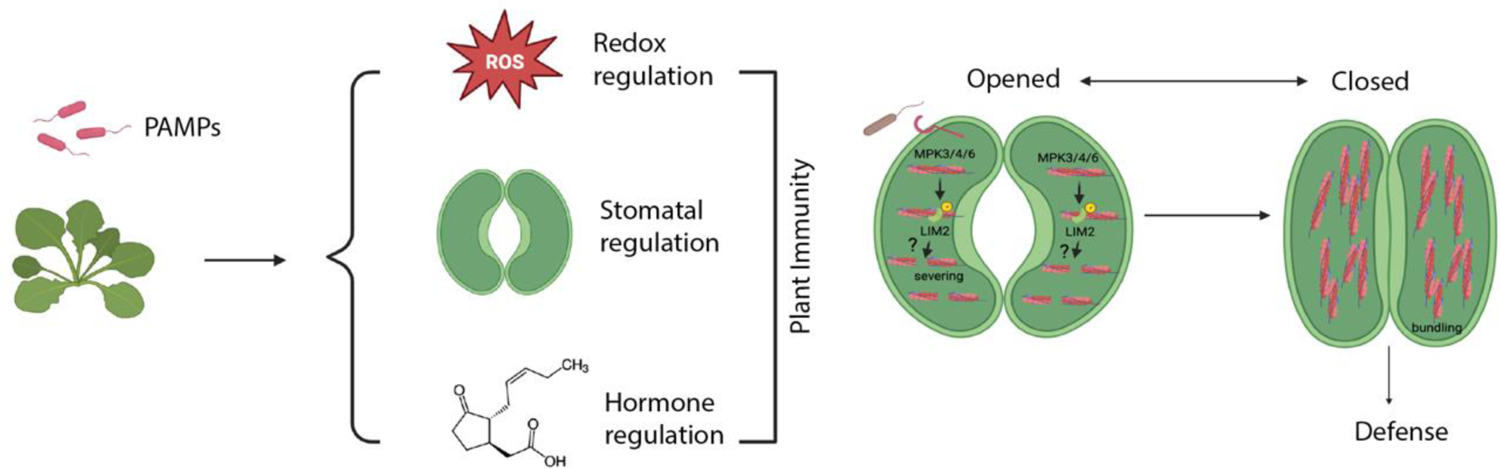
A working model for the role of WLIM2A in regulating actin dynamics during plant immunity. Upon MAMP perception, WLIM2A is phosphorylated by MPK3/4/6. This phosphorylation may stabilize the actin filaments, leading to initiate MAMP-induced PTI immune response. Upon pathogen infection, plants respond by redox changes, stomatal closure, and changes in the immune hormone levels. The WLIM2A protein may play an important role in redox regulation, stomata mediated immunity, and SA and JA biosynthetic pathways. Created with Biorender.com.

### Availability of data and material

RNA-Seq data are available at NCBI’s Gene Expression Omnibus GEO Series accession number GSE236403.

## Acknowledgments

We would like to thank the KAUST Bioscience Core labs for technical assistance for RNA sequencing and all members of the Hirt lab.

## Author contributions

PM, HH and NR designed the study. PM, AA, HA, MAT, and NR performed experimental work. PM, AV and NR performed *in silico* analysis and analyzed data. PM, AA, and NR wrote the paper. All authors read and approved the manuscript.

## Supplementary Material

**Figure S1.**
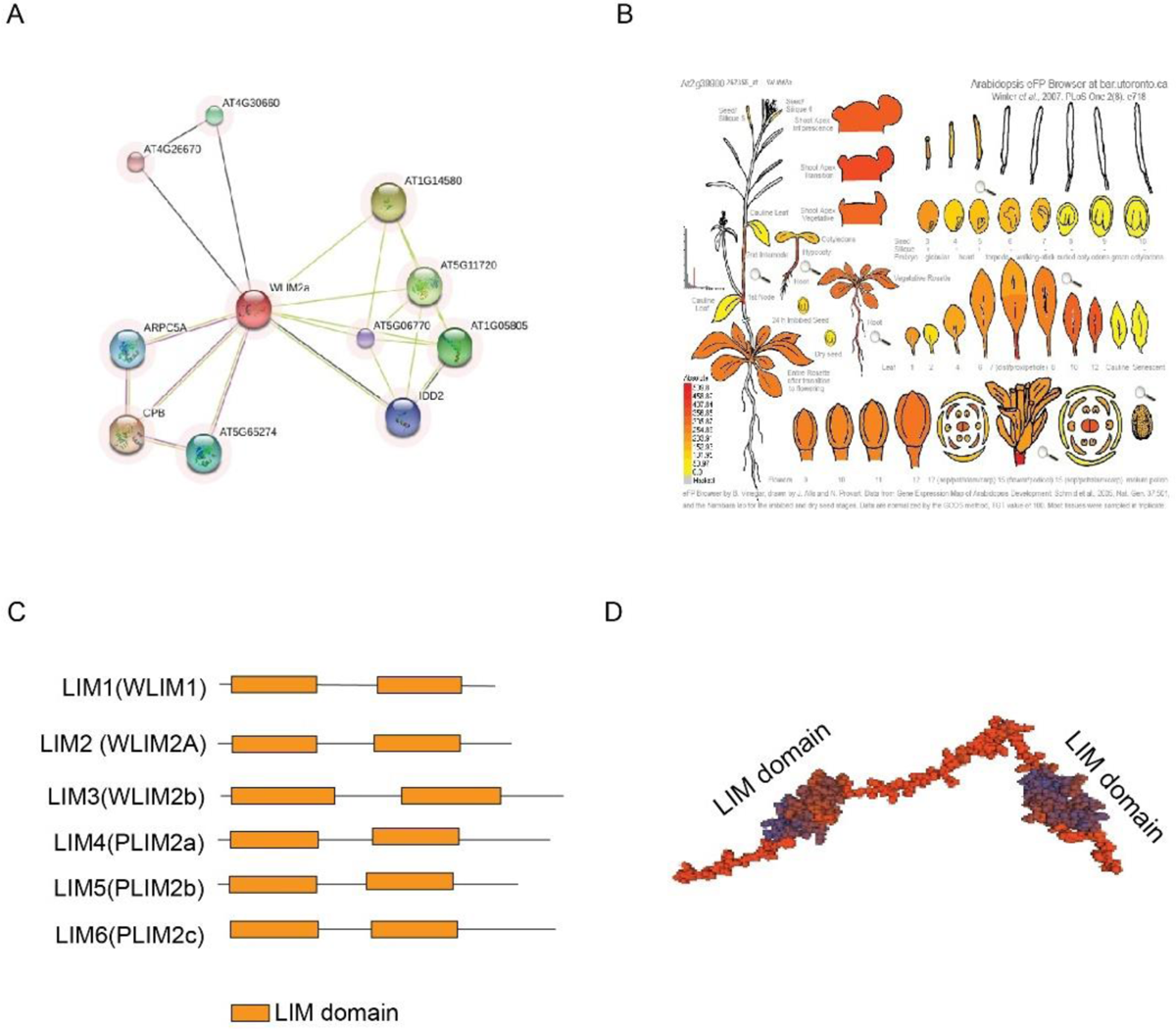
(**A**) Interaction networks from the STRING website, showing results of the potential partners for the WLIM2A protein. (**B**) eFP browser view of gene expression during *Arabidopsis* development. Expression of the WLIM2A, At2g39900, showing stronger expression widely seen in vegetative and reproductive stages. Expression strength coded by color: yellow=low, red=high. The *Arabidopsis* eFP Browser is located at bar.utoronto.ca, published in Winter et al.. 2007 (**C**) Schematic comparison of the domain architecture of the *AtLIM* gene family. (D) Predicted protein model of WLIM2A build from Swiss-model (Uniprot Id: O04193). The target sequence was performed against the SWISSMODEL template library. Two LIM domain contains double zinc finger motif at 8-68^th^ and 107-167^th^ position, respectively. Orange-full protein, blue and orange-LIM domain.

**Figure S2.**
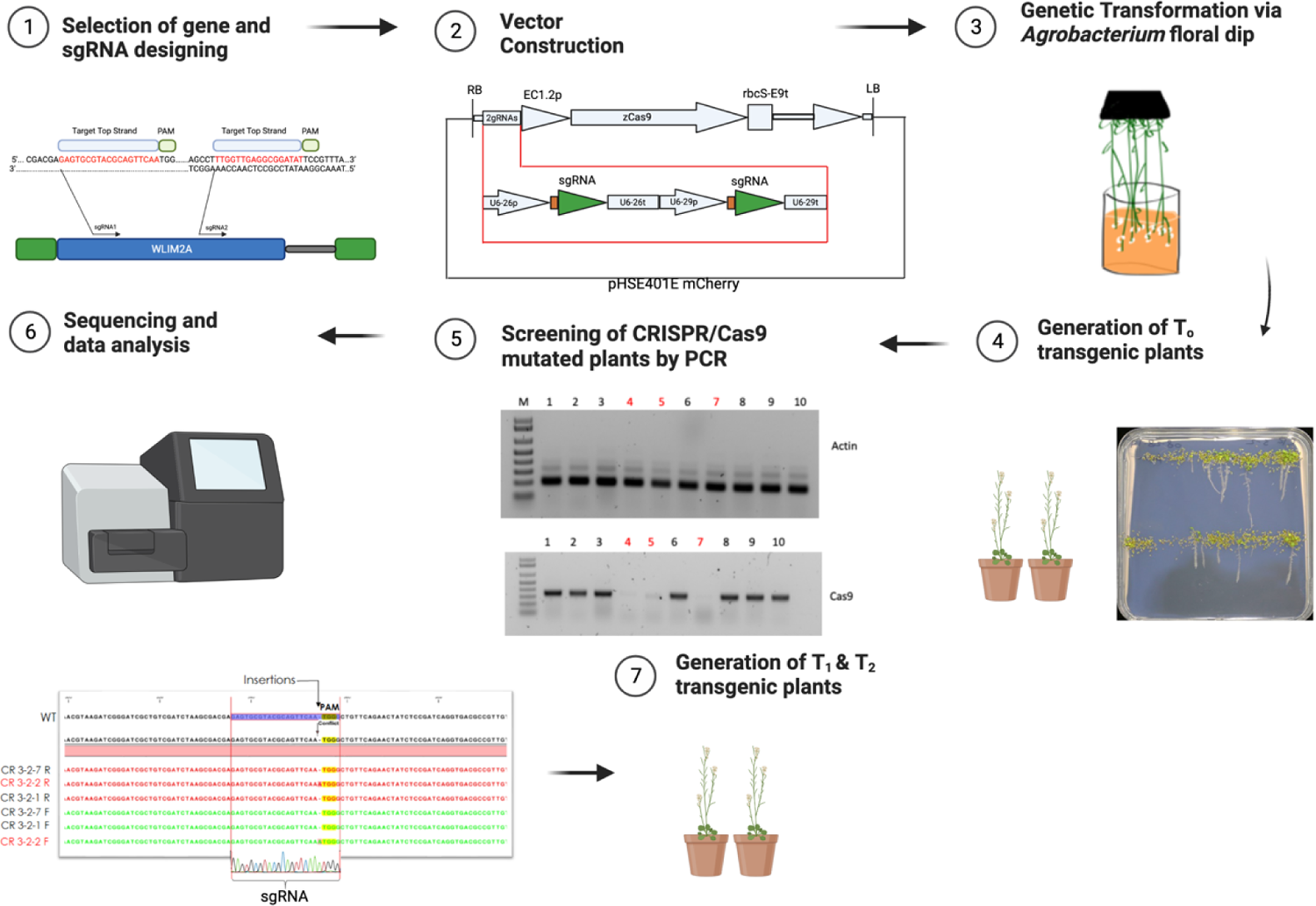
Workflow and Generation of CRISPR/Cas9 knockout mutant lines in *Arabidopsis*. Physical map of PHSE401 mcherry vector carrying two-gRNAs targeting two regions of *Arabidopsis* gene *WLIM2A.* The construct was transformed via *Agrobacterium* floral dip method. Screening of T1 transgenic plants on agar plates and selected the survival to grow in the soil. The DNA of plants was extracted, PCR amplified, and sequenced. Analysis of mutations using sequence alignment software to select for Indels and select homozygous mutants. Screening for T2 and T3 homozygous mutants and cas9 free lines is a key steps to generate CRISPR/Cas9 lines.

**Figure S3.**
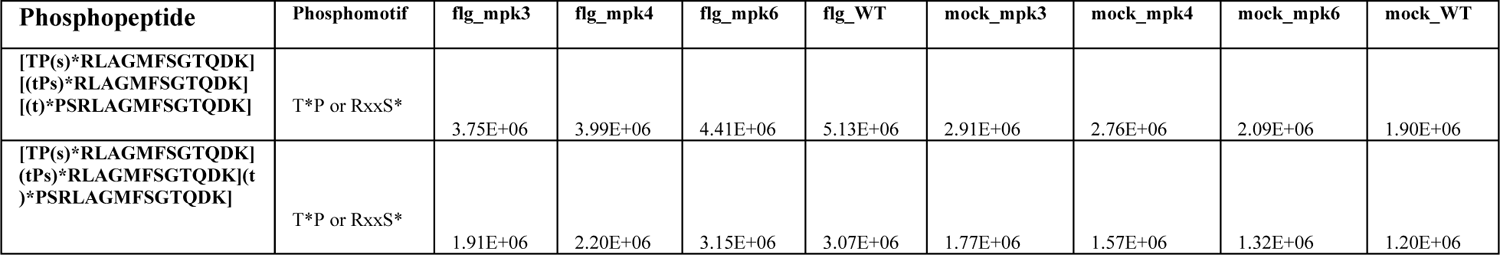
Table showing the relative abundance of the phosphopeptides. The abundance was measured after treatment by flg22 and/or the genotype (Rayapuram et al., 2018).

**Figure S4.**
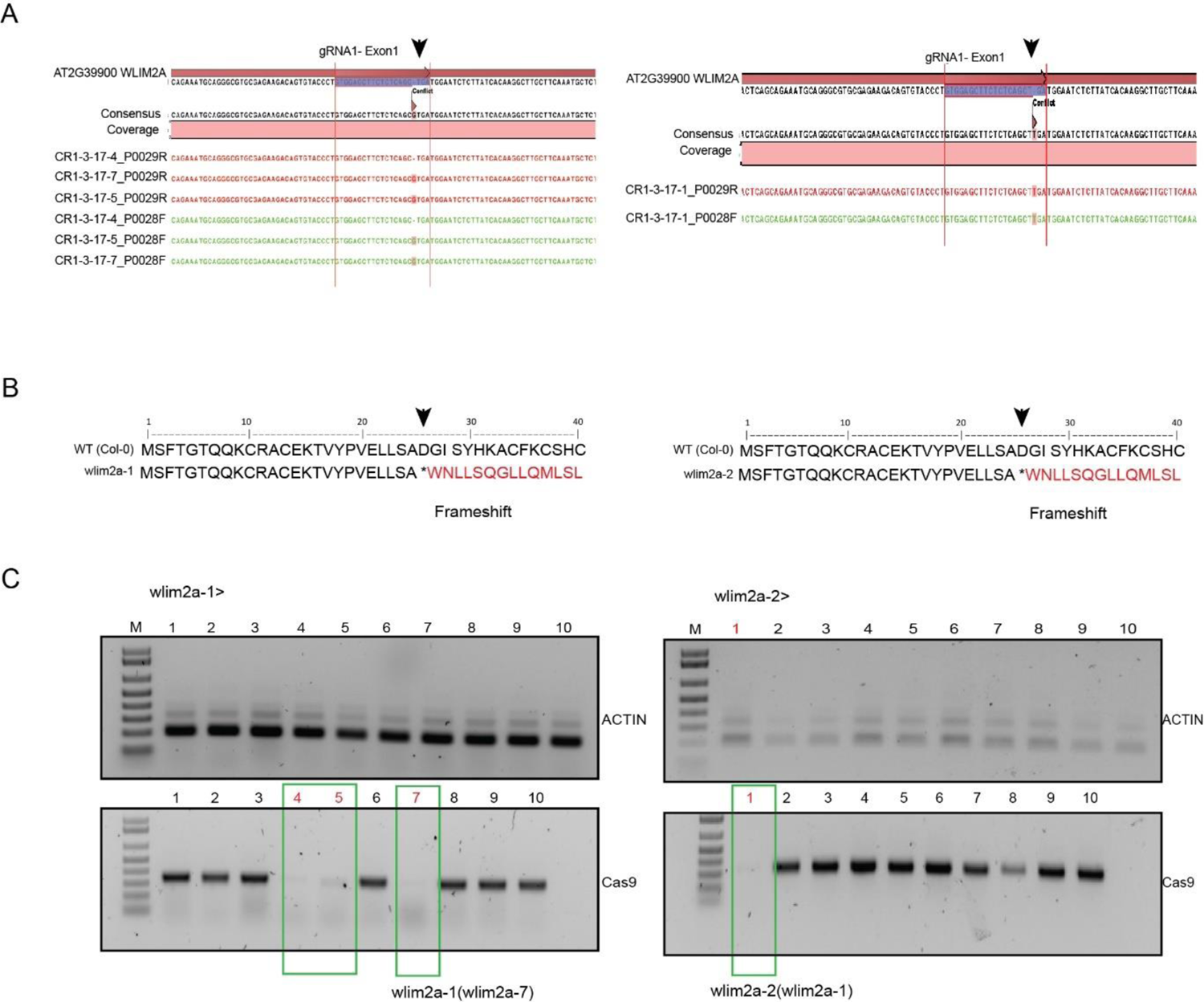
Inheritance of CRISPR/Cas9 mutations. (**A**) *In silico* analysis of nucleotide sequence of the targeted locus for Col-0 and *wlim2a*. One bp upstream of the PAM (TGA) within the sgRNA target sequence (highlighted), Cas9 cut site is indicated with a triangle (**B**) Sequence alignment of the targeted locus for Col-0, *wlim2a-1*, and *wlim2a-2*, all causing frameshift and formation of premature stop codon showing the frameshift indel resulting in a truncated protein of *wlim2a-1*, and *wlim2a-2.* (**C**) PCR amplification of Cas9 free CRISPR/Cas9 lines in T2 seedlings from the *wlim2a-1*(1-10), and *wlim2a-2* (1-10). Green boxed plants were Cas9 free lines.

**Figure S5.**
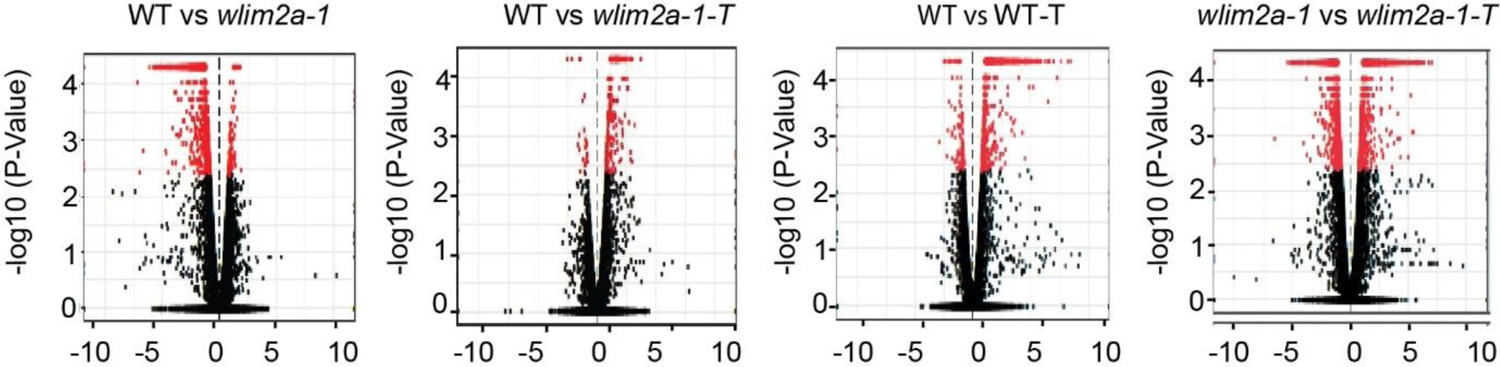
A comparison between mock and treated samples of WT and *wlim2a*-1 can be seen in the volcano plots. A plot is shown of log2 fold change against log10 (P-value).

**Figure S6.**
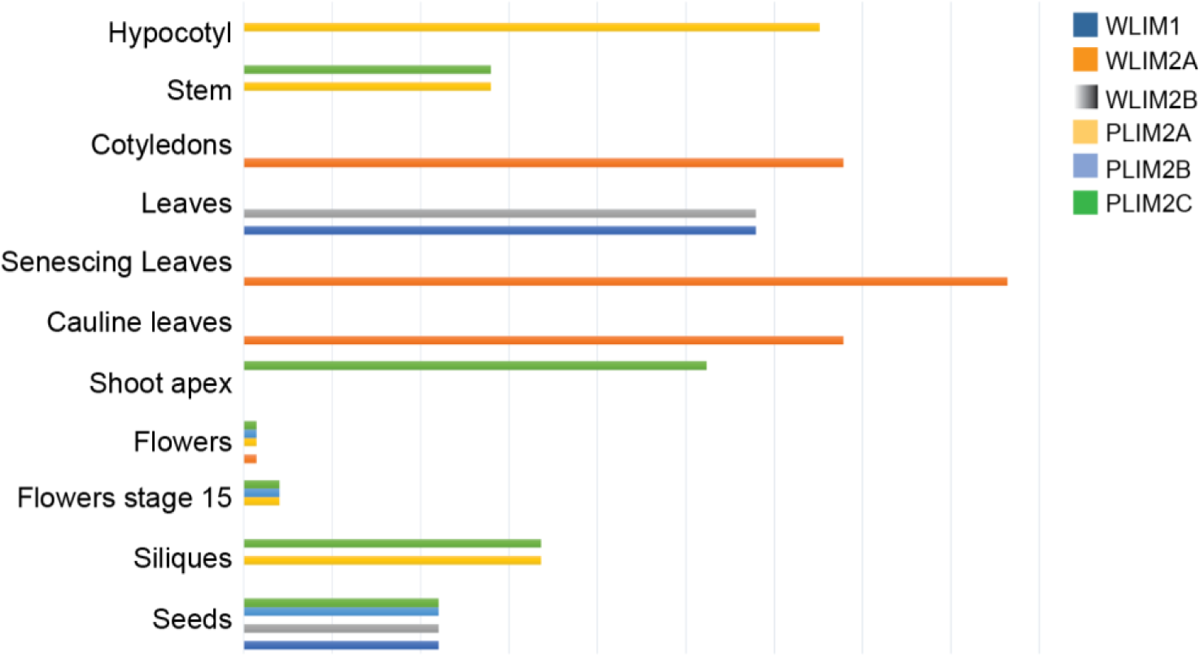
Microarray analysis of the LIM gene expression in *Arabidopsis*. (**A**) Electronic RNA gel blot analysis of LIM gene expression in different organs and development stages using the microarray. The data is retrieved from the AtGenExpress Schmid et al., 2005.

**Figure S7.**
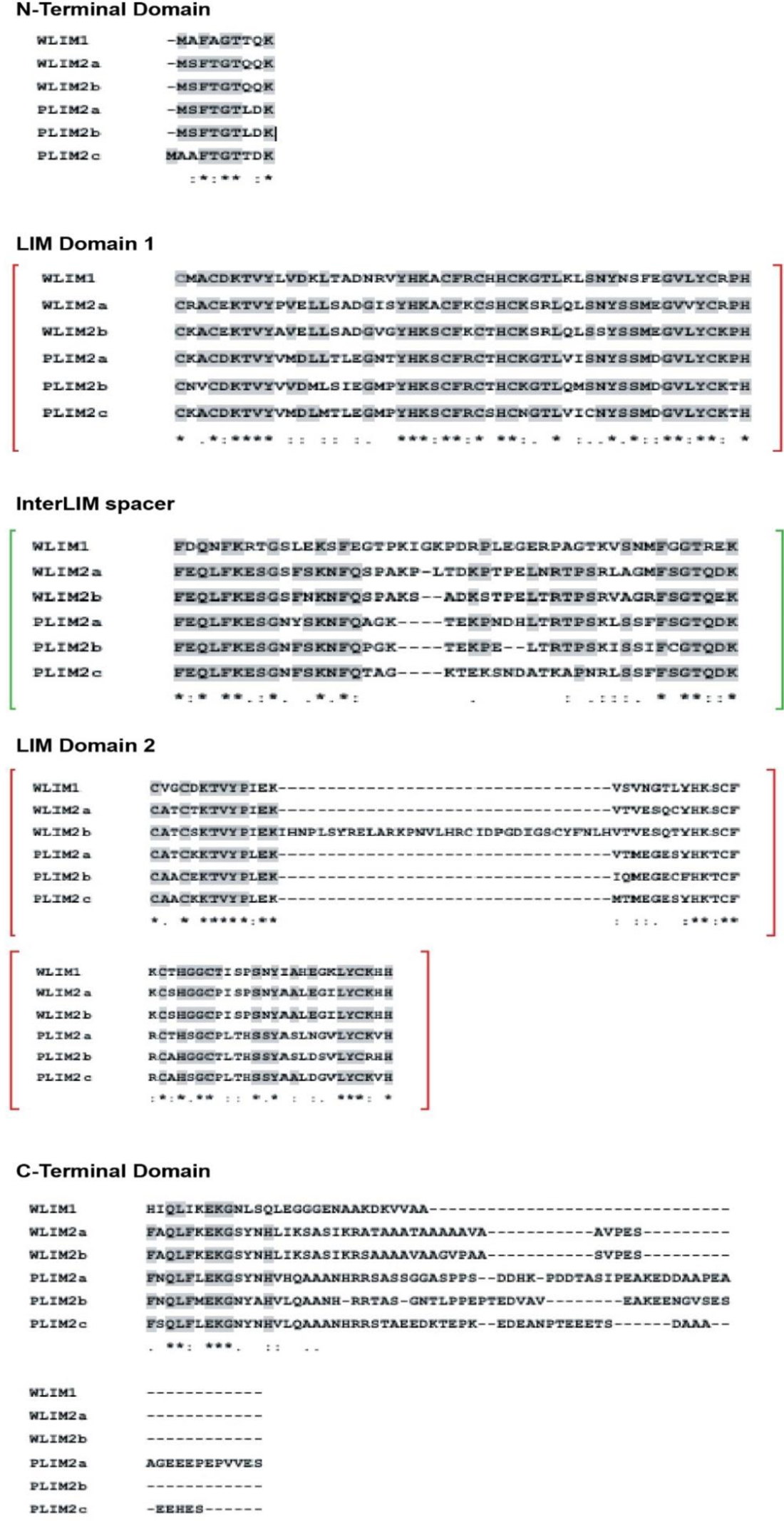
Multiple sequence alignment of *Arabidopsis* LIM proteins. Grey Shading indicates amino acids residues that are identical. The sequences were aligned using the Clustal Omega program(https://www.ebI.ac.uk/Tools/msa/clustalo/).The functional domains were identified through Uniprot (https://www.uniprot.org/uniprotkb/O04193/entry) and Pfam database (https://www.ebi.ac.uk/interpro/protein/UniProt/O04 193/entry/pfam/#table). The LIM domain and interLIMspacer are indicated in red and green, respectively.

